# Analysis of Moonlighting Proteins Regulated by Curcumin and Resveratrol in Cancer

**DOI:** 10.1101/2023.08.28.555039

**Authors:** Venil N. Sumantran, Anand Babu Kannadasan

## Abstract

Multitasking proteins or moonlighting proteins (MLP) play a major role in human disease and many are drug targets. MLPs have a single polypeptide with two different biochemical functions and often have different cellular localizations. Analysis of MultitaskProtDB-II and Comparative Toxicogenomics databases and Gene Ontology and STRING analysis, showed that curcumin and resveratrol regulate 3 groups of MLPs. Group 1 MLPs drive epithelial mesenchymal transition (EMT), whereas MLPs in groups 2 and 3 regulate tumour growth and suppression, respectively. The 3 groups of MLPs form a complex self-regulating network due to regulatory interactions between MLPs and the presence of transcription factors (CTNNB1, p53, NRF2) with their regulators and targets. Curcumin and/or resveratrol downregulate MLPs which drive EMT (CTNNB1, FASN, SMADs 2,3,4, MECP2, MMP2, TGFβR1, HES1) and promote growth and inflammation (MMP2, EGFR, FGF2, MDR1). Second, curcumin and/or resveratrol modulate pleiotropic, regulatory ‘hub’ MLPs (GSK3β, NOTCH1, HMGB1). Third, both phytochemicals upregulate tumour suppressor CDH1, and CYCS, which triggers apoptosis. Fourth, both phytochemicals modulate p53 expression, activity, and independently downregulate tumorigenic gene targets of p53 in groups 1 and 2. This dual mechanism can permit regulation of p53 activity in cells with wild type or mutant p53. Fifth, moonlighting functions of specific MLPs can cause unpredictable effects. These results add insights into synergistic and pleiotropic anti-cancer mechanisms of curcumin and resveratrol. It has clinical relevance because both phytochemicals are chemo-preventive agents and chemosensitizers in clinical trials.

## INTRODUCTION

Curcumin from turmeric and resveratrol from red wine are considered safe and beneficial food ingredients. Numerous preclinical studies show that curcumin can block carcinogenesis and inhibit tumour formation and progression in mouse models of colon, breast, lung cancers. These anti-cancer properties of curcumin require induction of growth arrest or apoptosis of premalignant and malignant cells. Accordingly, curcumin modulates key signalling molecules which regulate cell growth, survival, malignancy, and death [1,2]. Curcumin in combination with other phytonutrients is being evaluated as a food supplement for its chemo-preventive and immunomodulatory properties in lung and cervical cancer. Clinical trials are testing therapeutic potential of curcumin on localized cancers. Curcumin is also being tested as adjuvant with different chemotherapy agents, for therapeutic efficacy in adenocarcinomas and advanced cancers of the breast, colon, and rectum [2,3]. Various types of curcumin complexes and nano-curcumin formulations are also being tested for synergism with different chemotherapy agents in certain cancers [1, 2, 3]. Resveratrol targets many of the same molecules as curcumin, and has antiproliferative and antimetastatic activity in several cancer cell lines. Resveratrol also targets breast cancer stemlike cells and glioma stem cells from patient tissues [3]. Several derivatives and nano-formulations of curcumin and resveratrol have been developed since both phytochemicals have poor bio-availability [3,4]. Notably, curcumin formulations show improved delivery and anti-cancer activity in colon, breast, lung and liver cancer lines, and mouse models of colon and breast cancer [4].

Moonlighting proteins (MLPs) have a single polypeptide which performs two distinct biochemical functions in the same / different cellular locations. Moonlighting functions are often independent of the main function of a given MLP. Notably, almost 45% of MLPs are drug targets [5]. Curcumin and resveratrol have significant clinical potential and their mechanisms of action have been extensively studied in cancerous tissues [2,3]. However, there are no reports of MLPs regulated by both phytochemicals. Therefore, we identified all human MLPs regulated by curcumin and resveratrol. Gene Ontology (GO) and STRING analysis identified their functions and protein-protein interactions (PPI) respectively. Our results show that a subset of MLPs regulated by curcumin and resveratrol form a complex network of signaling molecules, transcription factors, oncogenes, tumor suppressors, and their regulators and targets. This MLP network regulates cell growth, death, malignancy, chemoresistance and immune resistance. We examined main and moonlighting functions of each MLP and its mode of regulation by curcumin and resveratrol. Our results provide novel insights into the tumorigenic impact of this MLP network, and the synergistic and pleiotropic anti-cancer mechanisms of these two phytochemicals.

## METHODS

### Identifying human Moonlighting Proteins Regulated by Curcumin and Resveratrol

The Comparative Toxicogenomics Database (CTD) [https://ctdbase.org/] curates relationships between thousands of chemicals, genes, phenotypes, and human diseases from the literature and other databases [Updated June 30, 2023. Revision 17133M]. The “Chemical-Gene Interaction” option in CTD identified 966 and 7254 curated proteins related to curcumin and resveratrol respectively. Venn analysis of these data identified 691 curated proteins related to both curcumin and resveratrol. A total of 684 curated moonlighting proteins (MLPs) were listed in the MultitaskProtDB-II database of moonlighting proteins, and 186 of these are human MLPs [5,6]. Venn analysis of 691 proteins related to curcumin and resveratrol with 186 human MLPs, identified 46 MLPs regulated by both curcumin and resveratrol. Figure 1 summarizes this workflow. Lists of 691 curated proteins related to curcumin and resveratrol, 186 human MLPs, and 46 MLPs regulated by curcumin and resveratrol, are provided in Supplementary Table 1.

**Figure 1:**
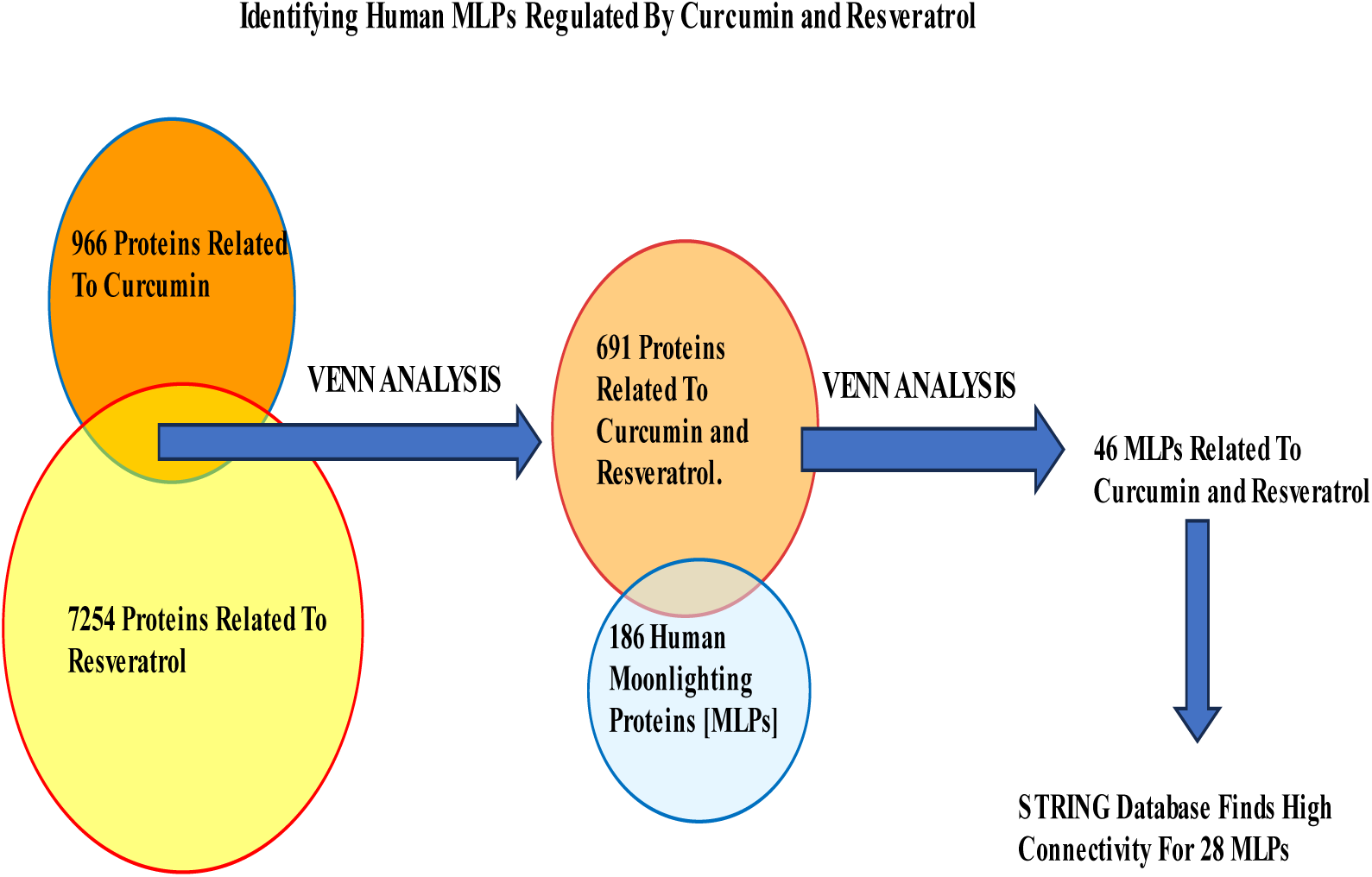
Identifying Human MLPs Regulated By Curcumin and Resveratrol. Comprehensive Toxicogenomic Database Identified Curated Proteins Related to Curcumin and Resveratrol. MultitaskProtDB-II identified Human MLPs. Venn Analysis identified 46 MLPs Regulated by both Curcumin and Resveratrol.

### Functions of human Moonlighting Proteins Regulated by Curcumin and Resveratrol

The Gene Cards database [https://www.genecards.org/ version 5.15] provided information on cellular localization and main molecular functions of the 46 moonlighting proteins regulated by curcumin and resveratrol. Moonlighting functions of MLPs were obtained from the Gene Cards database or the literature. Tables 2 and 3 summarize main and moonlighting functions of MLPs and their regulation by curcumin and resveratrol.

### Interactions of Moonlighting Proteins (MLPs) regulated by Curcumin and Resveratrol

Interactions between the 46 MLPs regulated by Curcumin and Resveratrol were visualized by STRING database which provides functional protein association networks [https://string-db.org/, STRING version 12.0]. STRING analysis was done with highest stringency [Confidence 0.90 and False Discovery Rate (FDR) of 1%]. Protein-protein interactions (PPI) with high scores from experimental evidence were used. Signalling Network Open Resource (SIGNOR) [http://signor.uniroma2.it/about/] confirmed these interactions between MLPs and provides scores for PPI based on causal interactions between human proteins, stimuli, phenotypes, enzyme inhibitors, complexes, and protein families [7]. We focussed on 28 of the 46 MLPs with highest PPI scores from STRING and SIGNOR databases. Interactions between 28 MLP with STRING and SIGNOR scores are in Supplementary Table 2.

### Regulation of Moonlighting Proteins (MLPs) by Curcumin and Resveratrol

Drug-gene interaction data from CTD listed how the 28 MLPs were regulated by curcumin and resveratrol. These data were verified with the cancer literature, and are in Tables 2 and 3. References for regulation of individual MLPs by curcumin and resveratrol are summarized in Supplementary Tables 3 and 4.

## RESULTS

### Gene Ontology (GO) and STRING Analysis of MLPs regulated by Curcumin and Resveratrol

The flowchart in Figure 1 summarizes how the 46 MLPs regulated by curcumin and resveratrol were identified. GO analysis showed that the 3 GO terms most significantly correlated with these 46 MLPs are; Regulation of Apoptotic process, Negative Regulation of Signal Transduction and Negative Regulation of Cell population Proliferation (Table 1).

**Table 1:** Gene Ontology (GO) terms correlated with 46 MLPs regulated by Curcumin and Resveratrol: Three GO terms most significantly correlated with Curcumin and Resveratrol are for Cell Death, Signalling, and Growth. Four MLPs (CTNNB1, TGFBR1, SMAD3, NOTCH1) are common to all 3 GO terms.

| Table 1 |  |  |  |  |
| --- | --- | --- | --- | --- |
| GO Term Number | GO Term | No. Genes | P-Value | Gene Symbols |
| GO 0042981 | Regulation of Apoptotic Process | 24 of 1550 | 4.99e-12 | **CTNNB1**, GSK3B, **TGFBR1**, **SMAD3**, **NOTCH1**, ATF2, HMGB1, MECP2, TIMP1, NME1, CRYAB, 90AB1, HSPD1, EGFR, CYCS, NOS3, P53, MDM2, PLK1, NRF2, NQO1, ENO1, XDH, GSN |
| <u>GO:0009968</u> | Negative Regulation of Signal Transduction | 15 of 1271 | 3.56e-06 | **CTNNB**, GSK3B, **TGFBR1**, SMAD 2, **SMAD 3**, SMAD 4, **NOTCH1**, FGF2, EGFR, NOS3, NRF2, P53, MDM2, ENO1, XDH, |
| <u>GO:0008285</u> | Negative Regulation of Cell population Proliferation | 12 of 696 | 2.25e-06 | **CTNNB1**, ATF2, **TGFBR1**, SMADs 2, **SMAD3**, SMAD 4, **NOTCH**, HES1, NME1, NOS3, P53, XDH |

**Table 2:**
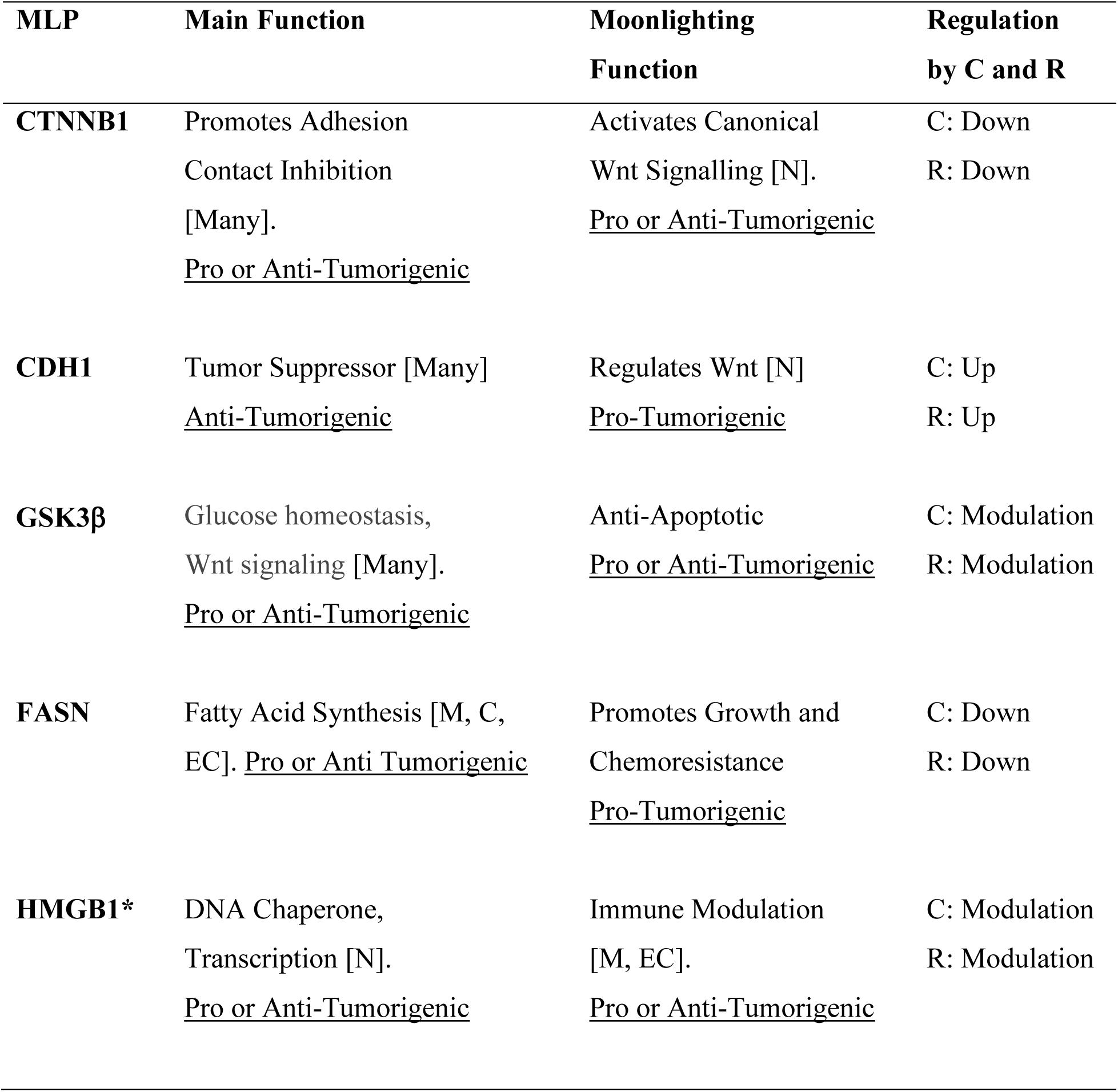
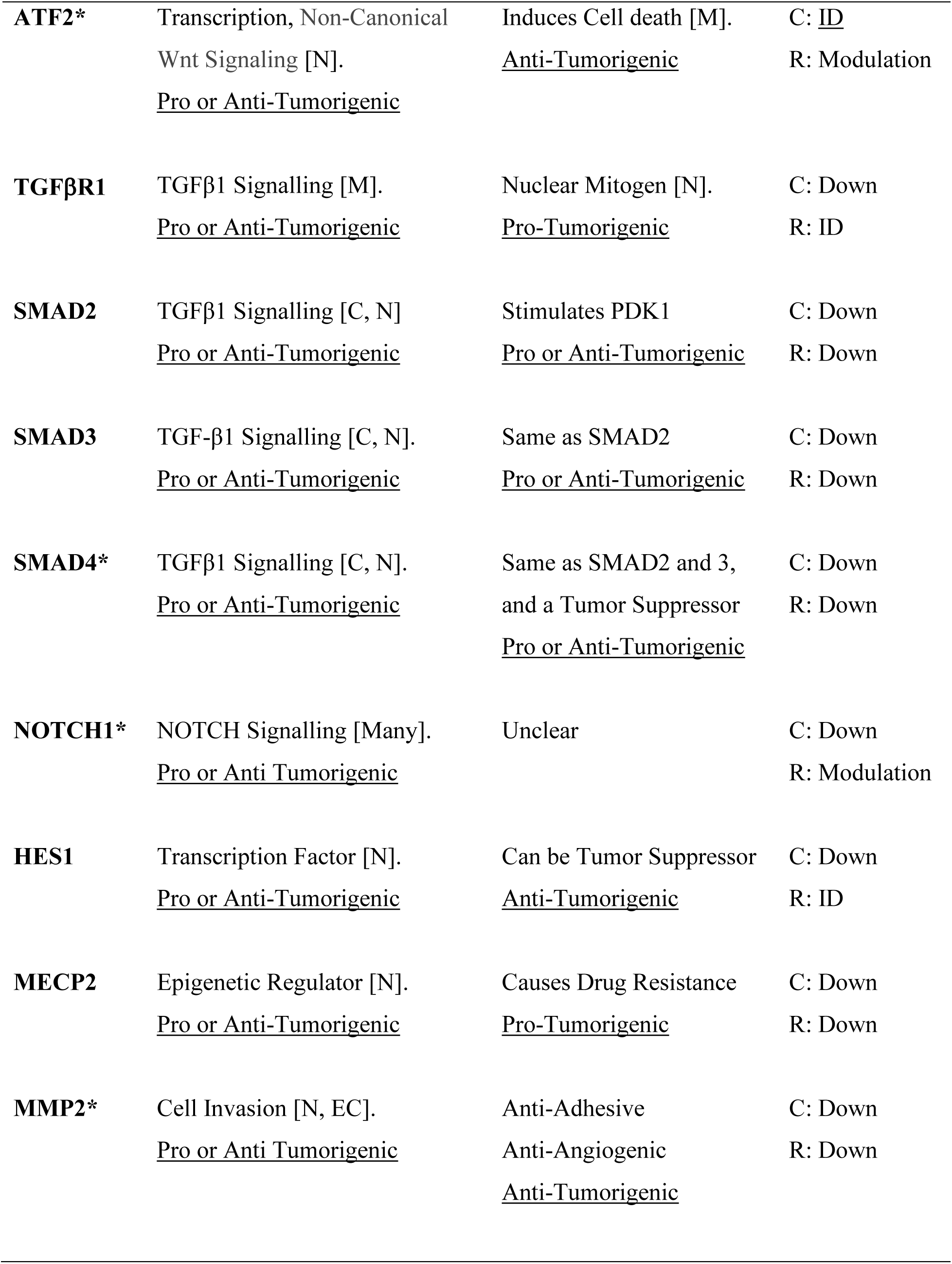
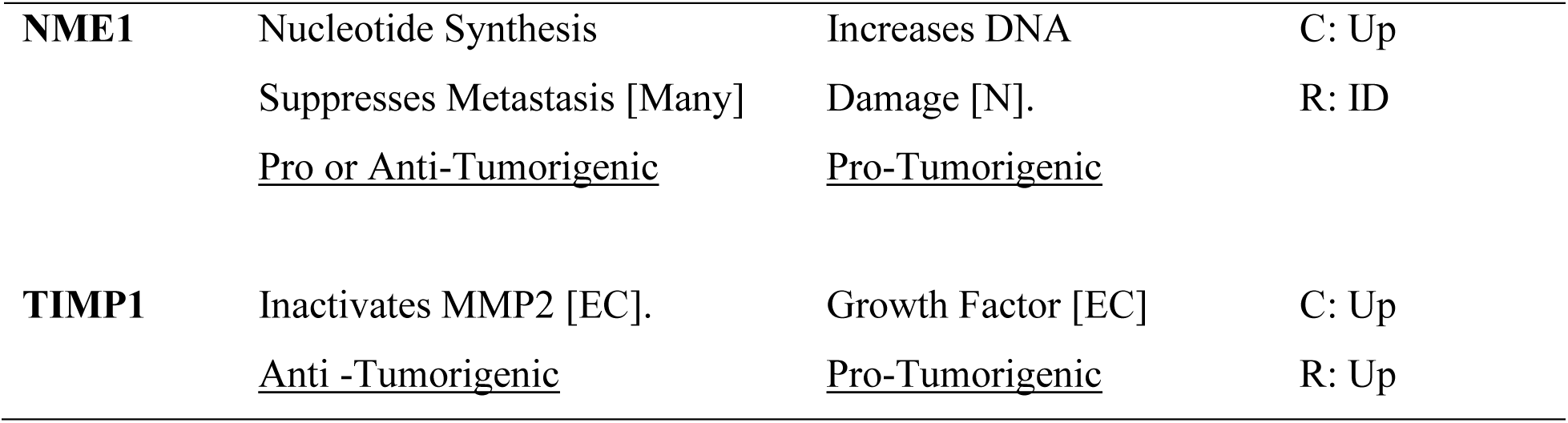
Regulation of Group 1 MLPs by Curcumin and Resveratrol (C and R). Four MLPs are Drug Targets (*). Upregulation (Up), Downregulation (Down), Modulation (Up OR Down). <u>Insufficient Data</u> (ID) for Some MLPs in Cancers. <u>Localization of MLP:</u> Nucleus (N), Cytoplasm (C), Plasma Membrane (M), Mitochondria (MT), Extracellular (EC). In 3 or more cellular compartments [Many].

These data are consistent with reports showing that anti-cancer mechanisms of curcumin and resveratrol primarily involve inhibition of cell growth and induction of growth arrest or cell death [1,2]. STRING analysis showed that 28 of the 46 MLPs regulated by Curcumin and Resveratrol have highest connectivity and PPI scores from both STRING and SIGNOR databases (Figure 2, Supplementary Table 2). Analysis of the STRING and GO terms showed that these 28 MLPs are in three functional groups. Group 1 contains sixteen MLPs which drive and modulate the process of epithelial mesenchymal transition (EMT). Four of these 16 MLPs (TGFβR1, SMAD3, NOTCH1, CTNNB1), are in all three GO terms of Table 1. These 4 MLPs regulate EMT and metastasis of tumour cells (GO:0010718: Positive regulation of epithelial to mesenchymal transition P=2.40e-07). Group 2 includes four heat shock proteins / chaperones, (CRYAB, HSP90AA1, HSP90AB1, HSPD1), and five pro-tumorigenic MLPs. CRYAB and HSPD1 can regulate apoptosis and are therefore in Table 1 (GO:0042981). Thus, group 2 has nine MLPs which regulate cellular stress and growth.

**Figure 2:**
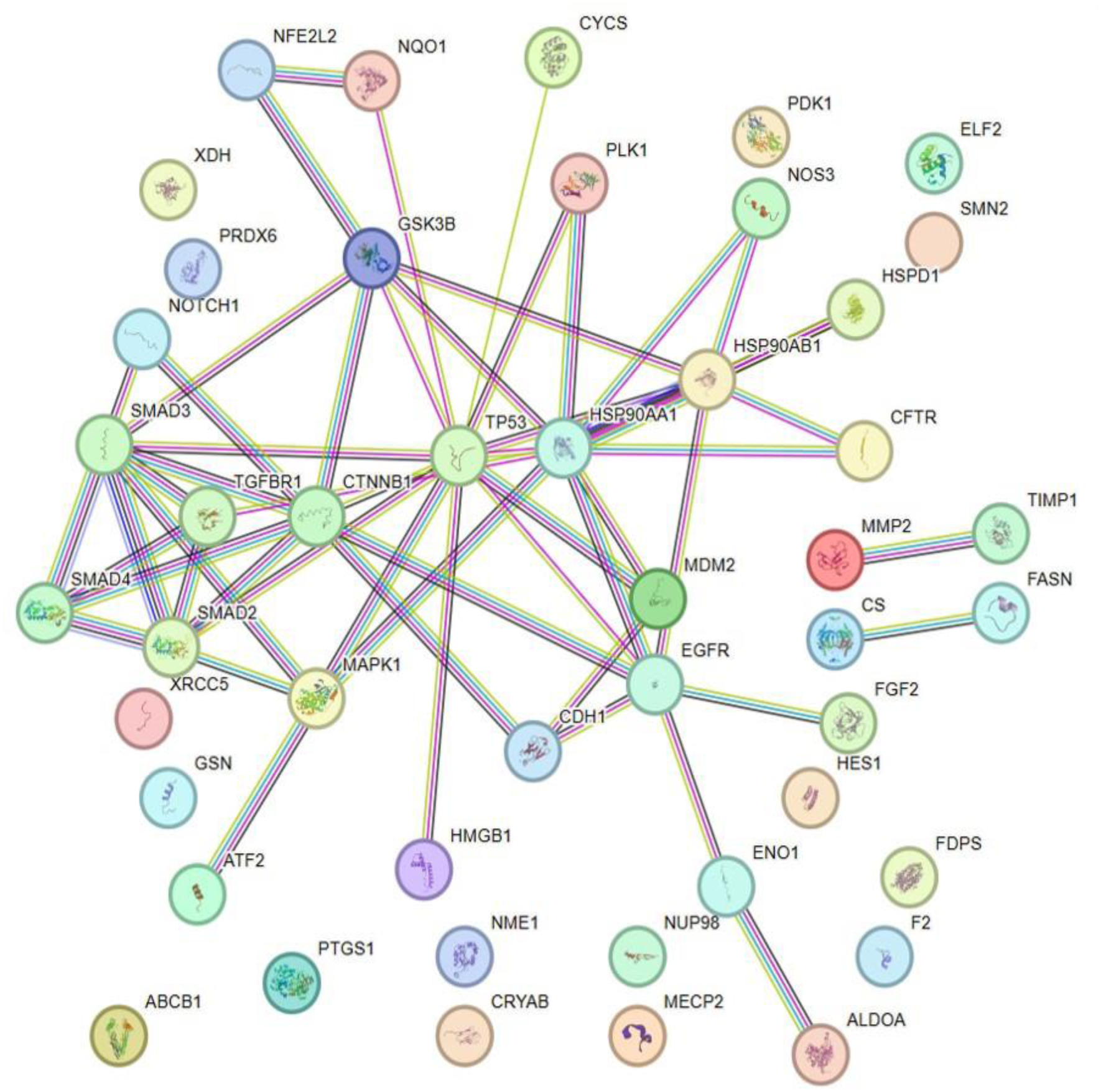
STRING Analysis of 46MLPs Regulated by Curcumin and Resveratrol. There are Significant Protein-Protein Interactions (PPI) between the 46 MLPs (Confidence 0.90, FDR (False Discovery Rate) = 1% and PPI Enrichment P-value = 4.44 e-16).

Group 3 has five MLPs including tumour suppressors, chemo-preventive MLPs, and their regulators and targets. Notably, all five MLPs in group 3 regulate cell survival and death (P53, MDM2, PLK1, NRF2, NQO1), and are therefore in Table 1 (GO:0042981).

In summary, GO and STRING analysis show that the 46 MLPs regulated by curcumin and resveratrol mainly regulate cell death, signaling and cell growth (Table 1). STRING data show statistically significant interactions between these MLPs (Figure 2). Interestingly, 28 of the 46 MLPs form a complex self-regulating network. These 28 MLPs are in 3 groups which regulate the EMT process, cellular stress and tumour growth, and tumour suppression, respectively (Figure 3). The following sections explain tumorigenic functions and interactions of each group of MLPs, and the pro or anti-tumorigenic effects of regulation of these MLPs by curcumin and resveratrol.

**Figure 3:**
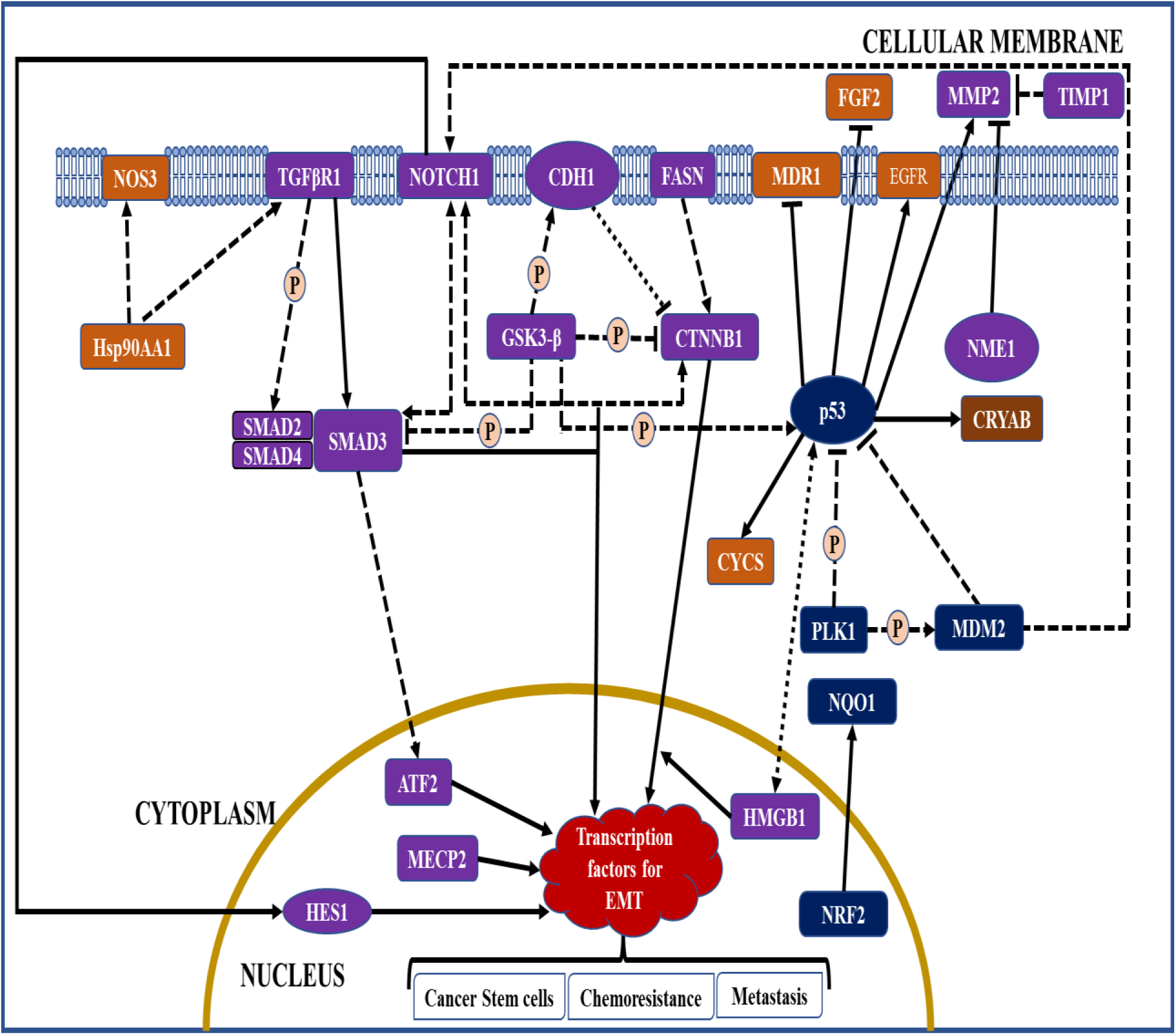
MLP Network Regulated by Curcumin and Resveratrol. Network contains 16 Group 1 MLPs 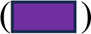, 7 Group 2 MLPs 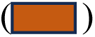, and 5 Group 3 MLPs 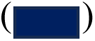. MLP interaction: Transcription regulation 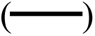, Protein-Protein Interaction 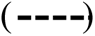, and complex formation 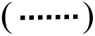, Positive regulation 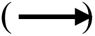, Negative regulation 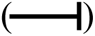.

### Group 1 MLPs regulate Epithelial to Mesenchymal transition (EMT)

The EMT process regulates wound healing and organ development in normal cells. However, EMT can become irreversible and drive tumour progression by converting epithelial cells into invasive, metastatic, mesenchymal cancer cells [8]. EMT also promotes increased resistance to chemo and radiotherapy agents, and cancer stem cell development [9]. The 16 MLPs in group 1 which regulate the EMT process are primarily in the Wnt, TGFβ1, and NOTCH1 pathways. Functions and interactions of each MLP, and their regulation by curcumin and resveratrol is explained.

### Group 1 MLPs in Wnt Pathway

CTNNB1, CDH1, and GSK3β are important MLPs in Wnt signalling. In the absence of Wnt ligands, the main function of CTNNB1 (β-catenin) is maintenance of cell adhesion and contact inhibition. However, in presence of Wnt ligands, CTNNB1 moonlights to activate transcription factors for canonical, β-catenin-dependent, Wnt signalling [10]. β-catenin is regulated by the moonlighting proteins CDH1 and GSK3β. CDH1 (E-cadherin) and CTNNB1 form complexes which maintain epithelial cell adhesion and survival, and block Wnt signalling by sequestering CTNNB1 in the membrane (Figure 3, Supplementary Table 2). Accordingly, CDH1 is a major tumour suppressor whose expression significantly correlates with improved survival of cancer patients [11]. CDH1 can moonlight to regulate Wnt signalling within nuclei [12] (Table 2). The main function of GSK3β (glycogen synthase kinase 3β) is to regulate glucose levels and activity of other proteins, but it moonlights to promote anti-apoptotic activity, independent of its kinase activity [13] (Table 2). Active GSK3β can inhibit Wnt signalling by two mechanisms. During absence of Wnt ligands, GSK3β can phosphorylate CTNNB1 and trigger its degradation. GSK3β can also phosphorylate CDH1 and promote its binding with CTNNB1, thus indirectly inhibiting Wnt signalling (Figure 3, Supplementary Table 2).

Increased chemoresistance and immune resistance are promoted by canonical Wnt signalling, and are mediated by FASN (fatty acid synthase) and HMGB1 (High Mobility Group B1) respectively (Figure 3). FASN is a multifunctional enzyme which moonlights to promote tumour cell growth and chemoresistance (Table 2). FASN can activate β-catenin / CTNNB1, and FASN inhibitors caused selective cell death in vivo xenograft tumour models by inhibiting β-catenin signalling and other pathways [14]. Table 2 shows that the main function of HMGB1 is to regulate organization, repair, and transcription of DNA. Notably, HMGB1 moonlights as an immunomodulator to protect normal cells, but it can enhance immune resistance in tumours [15,16]. Interestingly, HMGB1 and P53 form complexes which affect chemoresistance by regulating autophagy and apoptosis (Figure 3, Supplementary Table 2). The main functions of ATF2 (Activating Transcription Factor2) are pro or anti tumorigenic, but it can induce non-canonical, β-catenin independent, Wnt signalling in colon cancer [17]. Interestingly, ATF2 has an anti-cancer moonlighting function because it can promote apoptosis by altering mitochondrial permeability [18] (Table 2).

In terms of regulation, β-catenin / CTNNB1 is the crucial transcription factor for driving canonical Wnt signalling. Curcumin and resveratrol can inhibit expression and/or nuclear translocation of CTNNB1 in cancer cell lines, and resveratrol also downregulated CTNNB1 in colon cancer xenografts. GSK3β and CDH1 are negative regulators of CTNNB1. Both phytochemicals modulate (up/down regulate) GSK3β expression to cause apoptosis in cancer cell lines. The tumour suppressor CDH1, is upregulated by curcumin and resveratrol in cancer cell lines. Notably, curcumin could upregulate CDH1 in cancer stem cells and mouse colorectal tumours (Table 2 and Supplementary Table 3). FASN and HMGB1 mediate Wnt-dependent chemoresistance, and FASN was downregulated by curcumin and resveratrol in drug resistant and metastatic cancer cell lines (Table 2 and Supplementary Table 3). HMGB1 is novel because it’s upregulation and downregulation can be anti-tumorigenic. Thus, both phytochemicals can induce pyroptosis (immunogenic death) of tumour cells by stimulating HMGB1 secretion. However, curcumin and resveratrol also downregulate intracellular HMGB1 to induce apoptosis and decreases immune resistance of certain tumours. Little is known about effects of curcumin on ATF2 in cancers. However, resveratrol’s anti-carcinogenic effects are mediated by modulation of ATF2 (Table 2 and Supplementary Table 3).

### Group 1 MLPs in TGFβ1 Pathway

While Wnt ligands are expressed in epithelial and mesenchymal cells, TGFβ1 (transforming growth factor β1) is produced by many cell types and infiltrating immune cells after wounding or inflammation. TGFβ1 signalling is initiated by TGFβR1 (TGFβ1 receptor) which is pro or anti-tumorigenic because it regulates growth arrest, extracellular matrix production, immunosuppression, and carcinogenesis. Signals initiated by TGFβR1 are mediated by regulatory SMADs 2-4 (Sma- And Mad-Related transducer proteins 2,3,4), which translocate into the nucleus to induce expression of transcription factors required for EMT [8] (Figure 3). In early tumours, TGFβ1 is tumour suppressive because of its SMAD dependent, anti-proliferative, and pro-apoptotic effects. As tumours progress, SMAD independent effects cause TGFβ1 resistance and tumour promotion [19]. Interestingly, a nuclear fragment of TGFβR1 moonlights to promote mitosis and co-regulate transcription of SNAIL1, for the EMT process [20]. Therefore, TGFβR1 has a pro-tumorigenic moonlighting function. Independent of TGFβ1 ligand, SMADs 2,3,4 moonlight to stimulate PDK1 (Pyruvate Dehydrogenase Kinase 1), which can inhibit aerobic respiration, protect cells during hypoxia and oxidative stress, and inhibit TGFβ signalling [21]. Accordingly, moonlighting functions of SMADs 2,3,4 are pro or anti-tumorigenic. Notably, SMAD 4 can moonlight as tumour suppressor in certain cancers [22] (Table 2).

Regarding interactions, SMADs 3 and 4 crosstalk with GSK3β and ATF-2 in the Wnt pathway. Thus, GSK3β can phosphorylate and trigger degradation of SMAD3, which causes reduced sensitivity to TGFβ1. SMADs 3 and 4 can also activate ATF-2, which can activate non-canonical Wnt signalling [17], (Figure 3, Supplementary Table 2). In terms of regulation, curcumin decreases TGFβR1 expression and activity, whereas curcumin and resveratrol can decrease expression and phosphorylation of SMADs 2 and 3 in rats. Notably, resveratrol downregulated SMADs 2 and 3 in human breast cancer xenografts, and both phytochemicals downregulated SMAD4 in cancer cell lines (Table 2, Supplementary Table 3).

### Group 1 MLPs in NOTCH Pathway

Group 1 MLPs regulated by curcumin and resveratrol also include NOTCH 1 receptor and the HES1 (Hairy and Enhancer Of Split 1) transcription factor in the NOTCH signalling pathway. NOTCH signalling regulates growth, death, differentiation, and can stimulate expression of proteins linked with the EMT process and development of colon cancer stem cells [8,23]. HES1 can mediate NOTCH 1 induced EMT, and moonlight as a tumour suppressor in acute myeloid leukaemia [24, 25] (Table 2, Supplementary Table 2). Figure 3 shows that NOTCH1 can interact with SMAD3, CTNNB1, and MDM2. Thus, NOTCH1 and SMAD3 stabilize each other and activate expression of TGFβ1 dependent genes for EMT. Interactions between NOTCH1 and CTNNB1 modulate canonical Wnt signalling and NOTCH1 activity. MDM2 (Mouse Double Minute 2) usually degrades other proteins with its ubiquitin ligase activity, but MDM2 activates NOTCH 1 (Figure 3, Supplementary Table 2). Therefore, NOTCH 1 receptor can activate the EMT process via HES1 or by activating TGFβ1 or Wnt signalling. Studies in cancer cell lines show that curcumin inhibits NOTCH 1 expression and activity, whereas resveratrol modulates (up/down regulates) it. Curcumin also inhibits HES1 expression in various cancer cell lines (Table 2, Supplementary Table 3).

### Other Group 1 MLPs

In addition to MLPs in the Wnt, TGFβ1, and NOTCH 1 pathways, group 1 has four other MLPs which regulate the EMT process (MECP2, MMP2, TIMP1, NME1). MECP2 (Methyl-CpG-binding protein 2) is an epigenetic regulator of development which can promote EMT and metastasis, and it moonlights to increase multidrug resistance in some cancers [26,27] (Table 2). The second MLP is MMP2 (Matrix metalloproteinase 2), with main functions of regulating normal vascular development and wound healing. However, MMP2 can promote EMT by stimulating inflammation, angiogenesis, and cell invasion. Interestingly, MMP2 has an anti-tumorigenic moonlighting function because its C-terminus has a non-catalytic fragment (PEX) with anti-adhesive and anti-angiogenic properties [28] (Table 2). Notably, MMP2 can be regulated by the Wnt pathway and p53, in group 3. The third MLP which regulates the EMT process is NME1 / NM23 (Nucleoside diphosphate kinase 1), with a main function in pyrimidine synthesis, but NME1 also inhibits MMP2 and suppresses metastasis [29]. Interestingly, NME1 moonlights as an exonuclease to increase DNA damage [30]. Thus, NME1 has pro or anti-tumorigenic main functions and pro-tumorigenic moonlighting functions. The fourth MLP which regulates EMT is TIMP1 (tissue inhibitor of metalloproteinase 1). TIMP1 has anti-tumorigenic main functions because it irreversibly inactivates MMP2, and pro-tumorigenic moonlighting functions because it is mitogenic [31].

Overall, NME1 and TIMP1 can inhibit the EMT process by inhibiting MMP2 activity (Figure 3, Supplementary Table 2). With respect to regulation, curcumin and resveratrol downregulate MECP2 in cell lines and tissues of human breast cancer. Both phytochemicals inhibit expression and /or activity of MMP2, and upregulate TIMP1. Notably, curcumin inhibited tumour metastasis and upregulated both MMP2 inhibitors (NME1 and TIMP1) in mouse tumour models (Table 2, Supplementary Table 3).

To summarize, the 16 MLPs of group 1 drive the EMT process and are in the Wnt, TGFβ1, and NOTCH signalling pathways (Figure 3). Most of these MLPs have pro/anti-tumorigenic main functions. However, 8 MLPs (CTNNB1, FASN, TGFβR1, SMADs 2,3,4, MECP2, and MMP2) are pro-tumorigenic when overexpressed. Furthermore, FASN, TGFβR1, and MECP2, also have pro-tumorigenic moonlighting functions. Therefore, downregulation of these 8 group 1 MLPs by curcumin and resveratrol can have potent anti-cancer effects (Table 2). GSK3β and NOTCH1 are important regulatory hubs in group 1, and HMGB1 has pro or anti-tumorigenic effects in different cellular locations. Notably, both phytochemicals modulate GSK3β and HMGB1 expression, and resveratrol modulates NOTCH 1. Curcumin and resveratrol upregulate tumour suppressor CDH1, *in vitro* and in mouse colorectal tumours. Thus, curcumin and resveratrol downregulate, modulate, or upregulate specific group 1 MLPs to inhibit EMT in cancer cells and mouse models (Table 2, Supplementary Table 3).

### Group 2 MLPs Regulate Cellular Stress Responses

Group 2 MLPs regulated by curcumin and resveratrol include four heat shock proteins / chaperones which promote cell survival (CRYAB, HSPD1/HSP 60, HSP90AA1, and HSP90AB1). The main function of CRYAB (Crystallin Alpha B) is to prevent tissue damage. HSPD1 regulates mitochondrial proteins, whereas the main function of HSP90AA1 (Heat shock protein 90 Family A, Class A, Member 1) is to stabilize transcription factors and signalling proteins. Interestingly, these four chaperones can moonlight as immunomodulators [16]. HSP90AA1 has important interactions with 5 MLPs in the network [GSK3β, EGFR, P53, TGFβR1, and NOS3]. Thus, HSP90AA1 can bind and modulate GSK3β from group 1 [32], and stabilize EGFR and p53 of groups 2 and 3 respectively [33]. HSP90AA1 can also activate TGFβR1 and NOS3 (nitric oxide synthase) in groups 1 and 2 respectively (Figure 3, Supplementary Table 2). There is insufficient data on regulation for HSPD1, but curcumin was cytoprotective by preventing downregulation of CRYAB in rats (Table 3, Supplementary Table 4). Notably, HSP90AA1 which moonlights to increase immune resistance, was inhibited by curcumin in xenograft models of pancreatic cancer [34]. Studies show that NOS3 is upregulated and activated by resveratrol (Table 3, Supplementary Table 4).

**Table 3:** Regulation of Group 2, Group 3 MLPs by Curcumin and Resveratrol (C and R). Ten MLPs are drug targets (*). Upregulation (Up) Downregulation (Down), Modulation (Up OR Down). <u>Insufficient Data</u> (ID) for Some MLPs in Cancers. <u>Localization of MLP:</u> Nucleus (N), Cytoplasm (C), Plasma Membrane (M), Mitochondria (MT), Endoplasmic Reticulum (ER), Extracellular (EC). In 3 or more cellular compartments [Many].

| **MLP** | **Main Function** | **Moonlighting Function** | **Regulation by C and R** |
| --- | --- | --- | --- |
| **CRYAB** | Chaperone [C, N]<br><u>Pro or Anti-Tumorigenic</u> | Immune Infiltration.<br><u>Pro or Anti-Tumorigenic</u> | C: Up<br>R: ID |
| **HSP-90AA1\*** | Chaperone. [Many]<br><u>Pro or Anti-Tumorigenic</u> | Immuno-resistance [EC].<br><u>Pro-Tumorigenic</u> | C: Down<br>R: ID |
| **NOS3 \*** | Nitric oxide Synthesis [Many].<br><u>Pro or Anti-Tumorigenic</u> | Unclear | C: ID<br>R: Up |
| **COX1\*** | Prostanoid Synthesis. [ER, EC]<br><u>Pro or Anti-Tumorigenic</u> | Mitogen<br><u>Pro-Tumorigenic</u> | C: Down<br>R: Down |
| **EGFR\*** | EGF Signalling [M, EC ]<br><u>Pro or Anti-Tumorigenic</u> | Chemoresistance [N]<br><u>Pro-Tumorigenic</u> | C: Down<br>R: Down |
| **FGF2 \*** | Angiogenic Mitogen [C, N, EC]. <u>Pro or Anti-Tumorigenic</u> | Chemoresistance [N]<br><u>Pro-Tumorigenic</u> | C: Down<br>R: Down |
| **MDR1\*** | Transporter [N, M, EC].<br><u>Pro or Anti-Tumorigenic</u> | Regulates Membrane<br>Functions. [M]<br><u>Pro or Anti-Tumorigenic</u> | C: Down<br>R: Down |
| **CYCS\*** | Aerobic Respiration.[C, N]<br><u>Pro or Anti-Tumorigenic</u> | Triggers Apoptosis [MT]<br><u>Anti-Tumorigenic</u> | C: Up<br>R: Up |
| **P53 \*** | Transcription Factor [C, N]<br><u>Anti-Tumorigenic</u> | Induces Apoptosis, [MT]<br><u>Anti-Tumorigenic</u> | C: Modulation<br>R: Modulation |
| **MDM2** | Degrades p53 [Many]<br><u>Pro-Tumorigenic</u> | Increases Apoptosis [M]<br><u>Anti-Tumorigenic</u> | C: Down<br>R: Up |
| **PLK1 \*** | Regulates Mitosis [Many]<br><u>Pro or Anti-Tumorigenic</u> | Regulates Cell Invasion<br><u>Pro-Tumorigenic</u> | C: Down<br>R: Down |
| **NRF2** | Chemoprevention [Many]<br><u>Pro or Anti-Tumorigenic</u> | Anti-Inflammatory<br><u>Anti-Tumorigenic</u> [N] | C: Modulation<br>R: Modulation |
| **NQO1** | Detoxification [C]<br><u>Pro or Anti-Tumorigenic</u> | Stabilizes p53 and p73.<br><u>Anti-Tumorigenic</u> | C: Up<br>R: Up |

### Group 2 MLPs Regulate Growth, Angiogenesis

Group 2 MLPs also include COX1/PTGS1 (Cyclooxygenase-1), EGFR (Epidermal Growth Factor Receptor), and FGF2 (Basic Fibroblast Growth Factor 2), which promote inflammation, cell growth, and angiogenesis in normal cells, respectively (Figure 3, Table 3). However, COX1, EGFR, FGF2, are oncogenic when overexpressed. In terms of moonlighting functions, COX1 may promote tumour cell proliferation and nuclear forms of EGFR and FGF2 moonlight to promote chemoresistance. Thus, resistance to the EGFR inhibitor gefitinib correlated with increased expression of nuclear EGFR, and nuclear FGF2 regulates breast cancer chemoresistance [35, 36]. Another important group 2 MLP is ABCB1 (ATP binding cassette sub-family B Member 1) or MDR1/ P-Glycoprotein. MDR1 is a membrane transporter with efflux activity that causes chemoresistance, but it moonlights as a lipid translocase and regulates several membrane functions [37] (Table 3). Group 2 also includes CYCS (Cytochrome C-Somatic), which is essential for aerobic respiration, but CYCS moonlights to trigger apoptosis [38]. In terms of regulation, curcumin and/or resveratrol can downregulate expression or inhibit activity of COX1/PTGS1, EGFR, FGF2 in tumour models. Both phytochemicals inhibited MDR1 expression and/or activity in drug resistant human cancer cell lines, and induced apoptosis by upregulating or stimulating release of CYCS in human cancer cell lines (Table 3 and Supplementary Table 4).

To summarize, the MLPs in group 2 include four chaperones (CRYAB, HSPD1, HSP90AA1, and HSP90AB1) and four growth-promoting MLPs (COX1, EGFR, FGF2, MDR1), which have pro or anti-tumorigenic main functions, but are pro-tumorigenic when overexpressed. Furthermore, COX1, EGFR, and FGF2 also have pro-tumorigenic moonlighting functions. Therefore, downregulation of these MLPs by curcumin and resveratrol can have potent anti-cancer effects. Curcumin can also be anti-tumorigenic by upregulating the cytoprotective CRYAB, and downregulating HSP90AA1, which moonlights to increase immune resistance. CYCS which triggers apoptosis is upregulated by both phytochemicals (Table 3, Supplementary Table 4). Notably, many group 2 MLPs are transcriptional targets of p53, an MLP of group 3. Thus, CRYAB, COX1/PTGS1, EGFR, and CYCS are transcriptionally activated by p53, whereas FGF2 and MDR1 are suppressed by p53 (Figure 3, Supplementary Table 2).

### Group 3 MLPs are Tumour Suppressors

Group 3 MLPs regulated by curcumin and resveratrol includes the tumour suppressor p53, and its regulators. The P53 tumour suppressor protein activates transcription in response to stress, DNA damage, oncogene activation, and accordingly induces growth arrest or cell death. Independent of its transcriptional activity, P53 moonlights to alter mitochondrial permeability and induce apoptosis [39]. The GSK3β kinase can phosphorylate and activate P53 protein, but MDM2 and PLK1 inhibit P53 (Figure 3, Supplementary Table 2). Thus, MDM2 ubiquitin ligase degrades P53 protein, and accordingly has pro-tumorigenic main functions (Table 3). Independent of its enzyme activity, MDM2 moonlights to regulate NDUFS1, decrease mitochondrial respiration, and increase apoptosis [40]. Thus, MDM2 has anti-tumorigenic moonlighting functions. PLK1 (Polo Like Kinase-1) regulates mitosis as its main function, and may moonlight to regulate vimentin and breast cancer cell invasion [41]. PLK1 inhibits P53 protein via two mechanisms. PLK1 can phosphorylate P53 and inhibit its transcriptional activity. PLK1 can also phosphorylate and activate MDM2, which then degrades P53 (Figure 3, Supplementary Table 2).

In terms of regulation, several reports show that curcumin and resveratrol upregulate expression of p53 and its target genes, which then induce growth arrest or apoptosis in cancer cells and some mouse models of cancer. However, curcumin and resveratrol can also downregulate p53 expression to prevent cell death and organ damage (Table 3, Supplementary Table 4). Accordingly, both phytochemicals can modulate (up/down regulate) p53 levels and cause cell death and cyto-protection respectively. Curcumin and resveratrol also modulate GSK3β, an activator of p53, and curcumin can downregulate both inhibitors of p53 (MDM2 and PLK1), and restore p53 activity. In contrast, resveratrol upregulates MDM2 and downregulates PLK1, to cause growth inhibition or cell death in different cancer cell lines (Table 3, Supplementary Table 4).

### Group 3 MLPs are Chemo-preventive

Group 3 also includes the chemo-preventive transcription factor NRF2 / NFE2L2 (NFE2 Like BZIP transcription factor 2) and its gene target NQO1 (NAD(P)H Quinone Dehydrogenase 1). While p53 protects from genotoxic damage, the main function of the NFE2L2 / NRF2 transcription factor is to upregulate antioxidant and cytoprotective genes such as superoxide dismutase, glutathione transferase, and NQO1 (Supplementary Table 2). NRF2 is chemo-preventive in normal cells but can also increase survival and chemoresistance of tumour cells. The main detoxification function of NQO1 also supports normal and tumour cells.

Accordingly, both NRF2 and NQO1 can have pro or anti-tumorigenic main functions. Independent of transactivating cytoprotective genes, NRF2 moonlights to prevent upregulation of pro-inflammatory cytokines by macrophages *in vitro* and in mice. [42]. During oxidative stress, NQO1 moonlights to prevent degradation of p53 via ubiquitin independent mechanisms [43]. Therefore, both NRF2 and NQO1 have anti-tumorigenic moonlighting functions (Table 3). In terms of regulation, the upregulation of NRF2 by resveratrol was chemo-preventive in animal models of carcinogenesis. Activation of NRF2 by curcumin inhibited EMT or prevented toxicity. However, inhibition of NRF2 nuclear translocation or activity by curcumin and / or resveratrol, reversed chemoresistance in different cancers. Therefore, both phytochemicals can have anti-cancer effects by modulating expression or activity of NRF2. Curcumin decreased liver toxicity by upregulating NQO1 (Table 3, Supplementary Table 4).

To summarize, group 3 MLPs include p53 and its regulators. Curcumin and resveratrol control the expression, stability, and activity of p53 by regulating expression of the activator (GSK3β), and inhibitors (MDM2 and PLK1) of p53. In cancer cells, both phytochemicals often upregulate p53 and its tumor suppressive function. However, downregulation of p53 can be cytoprotective. Thus, curcumin and resveratrol modulate (up/down regulate) expression of wild type p53 in different contexts. When curcumin and resveratrol upregulate or downregulate wild type p53, its pro-tumorigenic gene targets (MMP2, EGFR, FGF2, MDR1) can be indirectly upregulated or downregulated, respectively. However, both phytochemicals can also directly downregulate these same pro-tumorigenic gene targets of wild type p53. Thus, curcumin and resveratrol can directly downregulate MMP2, EGFR, FGF2, MDR1 in a cancer cell with wild type or mutant p53. Similarly, curcumin and resveratrol can indirectly or directly regulate expression of CYCS, which is another target of wild type p53. This dual mechanism of indirect and direct regulation of p53 gene targets is a novel anti-cancer mechanism by which both phytochemicals could target tumours irrespective of their p53 status. The cytoprotective NRF2 and its target NQO1, are also in group 3. Curcumin and resveratrol are anti-tumorigenic by modulating (up/down regulating) NRF2 in different contexts. Both phytochemicals can protect against drug toxicity by upregulating NQO1.

### Regulation of the 3 Groups of MLPs by Curcumin and Resveratrol

The venn diagram in Figure 4 summarizes how curcumin and resveratrol regulate the MLP network shown in Figure 3. Figure 4A shows that curcumin and resveratrol downregulate group 1 MLPs which drive EMT (CTNNB1, FASN, SMADs 2,3,4, MECP2, MMP2). In group 2, curcumin and/or resveratrol downregulate MLPs which promote growth, inflammation, and chemoresistance (COX1/PTGS1, EGFR, FGF2, MDR1). Interestingly, curcumin downregulates both inhibitors of p53 in group 3 (MDM2 and PLK1). Notably, GSK3β, NOTCH 1, and HMGB1 are important regulatory group 1 MLPs which are anti-tumorigenic when up / down regulated in different contexts. Figure 4B shows that these 3 MLPs can be modulated by curcumin and / or resveratrol. In group 3, p53 is a tumour suppressor and NRF2 is chemo-preventive, but can promote chemoresistance when overexpressed. Accordingly, both phytochemicals modulate p53 and NRF2 to give anti-cancer effects (Figure 4B, Supplementary Table 4). Figure 4C shows that curcumin and resveratrol upregulate the anti-tumorigenic CDH1 and TIMP1 in group 1 and CYCS in group 2. To summarize, the anti-cancer effects of curcumin and resveratrol involve very similar modes of regulation of specific MLPs in the 3 groups. Only NOTCH 1 and MDM2 are differentially regulated by curcumin versus resveratrol (Tables 2,3, Figure 4).

**Figure 4:**
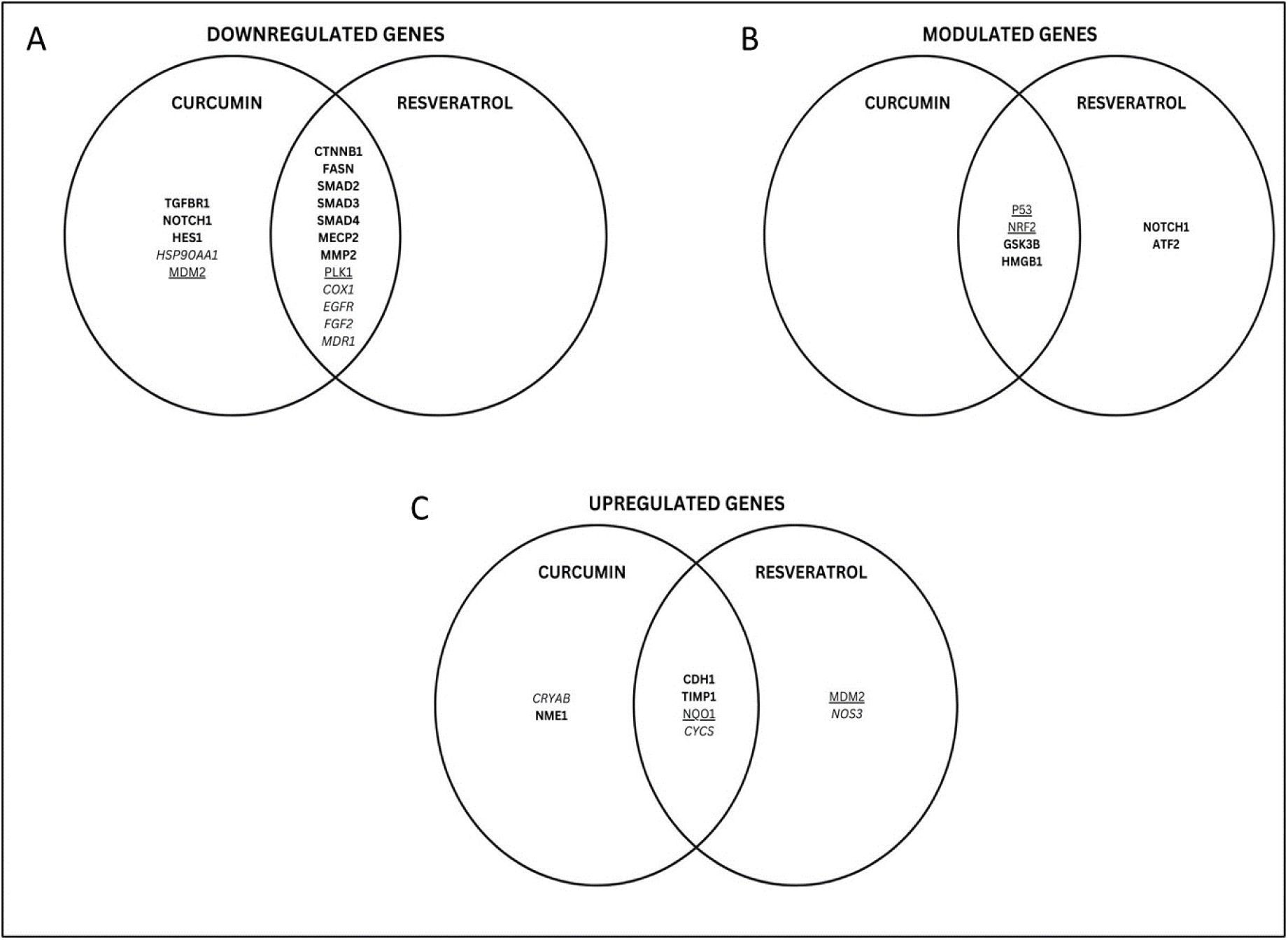
MLPs Regulated by Curcumin and Resveratrol. Downregulated MLPs (**4A**), Modulated MLPs (**4B**), and Upregulated MLPs (**4C**) Group 1 MLPs (Bold), Group 2 MLPs (Italics), and Group 3 MLPs (Underlined).

## DISCUSSION

The MLPs we analysed are important in cancer [46], and are known targets of curcumin and/or resveratrol [1–4]. Most studies focussed on main functions of these MLPs but did not analyse their moonlighting functions and their role in a complex, self-regulating network.

We explain how curcumin and resveratrol control this network of human MLPs which drive tumorigenesis and chemoprevention (Figure 3). Group 1 MLPs in the Wnt, TGFβ1, and NOTCH pathways, drive EMT, metastasis and chemoresistance. Notably, Group 2 MLPs regulated by curcumin and resveratrol include oncoproteins which are established drug targets [5]. Group 3 MLPs include the tumour suppressor p53, and MLPs which control stability and activity of p53. The chemo-preventive NRF2 and its target NQO1, are also in group 3. This network of three groups of MLPs is complex and self-regulated for three reasons. First, many group 1 MLPs regulate each other at the post-translational level. Second, many group 2 MLPs are transcriptional targets of p53, from group 3. Third, the network has activators and inhibitors of important MLPs. These include, CTNNB1 and its activators FASN and NOTCH1, ATF2 and its activator SMAD3, NOTCH 1 and its activator MDM2, and p53 and its activator GSK3β. The network also has inhibitors of CTNNB1, SMAD3, MMP2, and p53 (Figure 3). Regarding regulation, curcumin and resveratrol can modulate activities of GSK3β, SMADs 2,3 and inhibit COX1 and MDR1 activity. However, both phytochemicals primarily regulate MLPs at the transcriptional level (Supplementary Tables 3,4). Thus, curcumin and/or resveratrol downregulate MLPs which drive EMT, modulate regulatory MLPs (GSK3β, HMGB1, NOTCH1) and transcription factors (p53, NRF2), and upregulate tumor suppressor CDH1, detoxifier NQO1, and apoptosis inducer CYCS (Figure 3). These similarities in their regulation of MLPs provide a novel mechanism for synergistic anti-cancer effects of curcumin and resveratrol. These results agree with reports of synergism between curcumin and resveratrol in rodent models of prostatic and breast cancer [44,45].

Limitations of this study include insufficient information on moonlighting functions for 2 of the 28 MLPs (NOTCH1, NOS3) in cancers. There is also insufficient literature on regulation of ATF2 and NOS3 by curcumin, and regulation of TGFβR1, HES1, CRYAB and HSP90AA1 by resveratrol, in cancers (Tables 2,3). However, the remaining data provide five important insights about the MLPs regulated by curcumin and resveratrol. First, curcumin and resveratrol have novel, coordinated, anti-cancer mechanisms of regulating major MLPs such as CTNNB1, GSK3β, MMP2, p53, their regulators, and targets (Figures 3,4). Thus, CTNNB1 which drives EMT, is downregulated by curcumin and resveratrol, However, both phytochemicals can also downregulate FASN, the activator of CTNNB1, and upregulate CDH1, an inhibitor of CTNNB1. Another example is pro-tumorigenic MMP2, which is inhibited by both phytochemicals. However, curcumin also upregulated both MMP2 inhibitors (TIMP1 and NME1) in mouse tumours. The best example is the ‘dual’ mechanism by which curcumin and resveratrol regulate expression of p53, its activator (GSK3β), and inhibitors (MDM2, PLK1), but can also directly downregulate pro-tumorigenic gene targets of p53 in groups 1 and 2 (Figures 3,4). Examining main versus moonlighting functions of MLPs also provides insights. Thus, the second insight relates to MLPs which moonlight to modulate immune responsiveness (HSP90AA1 HSP90AB1, CRYAB, HMGB1, NRF2).

Moonlighting functions of these MLPs and their regulation has clinical relevance, because immune checkpoint inhibitor therapy is a promising new cancer treatment. Group 1 MLPs in Wnt pathway are also notable since Wnt signalling can regulate antitumor immunity and immune evasion [3]. Therefore, inhibition of Wnt signalling by curcumin and resveratrol may reduce immune resistance in some cancers. Third, several MLPs regulated by curcumin and resveratrol moonlight as regulators of cell death (GSK3β, CYCS, P53, MDM2, NQO1), cell growth (TGFβR1, TIMP1, COX1), and chemo-resistant growth (FASN, EGFR, FGF2, MECP2, NRF2). Notably, curcumin and resveratrol can trigger cell death by regulating moonlighting functions of HMGB1, P53, CYCS (Tables 2,3, and Supplementary Tables 3,4). However, studies focusing on the effects of curcumin and resveratrol on moonlighting functions of the remaining MLPs, will help us understand their regulation of cell death, growth, and immune resistance in cancers.

Examining the tumorigenic impact of the main versus moonlighting functions of MLPs also provides important insights. Thus, our fourth insight is if both phytochemicals regulate MLPs with opposing main and moonlighting functions (MMP2, TIMP1 and CDH1), it can cause pleiotropic or unpredictable ‘off-target’ effects which weaken inhibition of EMT [8]. Thus, MMP2 has pro-tumorigenic main functions when overexpressed, and anti-tumorigenic moonlighting functions. Curcumin and resveratrol downregulate MMP2, which could decrease its main and moonlighting functions, and weaken the expected anti-tumorigenic effects (Table 2). Similarly, curcumin and resveratrol upregulate the main anti-tumorigenic functions of TIMP1 and CDH, but may also upregulate their pro-tumorigenic moonlighting functions (Table 2), and weaken the expected anti-cancer effects. MDM2 also has opposing main and moonlighting functions, and its regulation by curcumin and resveratrol may cause pleiotropic effects on cell survival (Table 3). The fifth insight is that pleiotropic effects can also occur when curcumin and resveratrol regulate MLPs with main and moonlighting functions in different cellular locations. Five group 1 and 2 MLPs (CDH1, COX1, TGFBR1, EGFR, FGF2) have this property. Little is known on regulation of moonlighting functions CDH1 and COX1. However, tumorigenic impact of main and moonlighting functions of TGFβR1, EGFR, and FGF2 is reported [46]. Thus, a fragment of TGFβRI enters nuclei, moonlights as a co-transcriptional regulator of p300, and upregulates the SNAIL and MMP2 genes required for EMT [20]. Nuclear forms of EGFR and FGF2 moonlight to increase chemoresistance [35,36]. Notably, EGFR inhibitors for cancer patients only target membrane bound EGFR which performs the main functions of EGFR

## CONCLUSION

Curcumin and resveratrol regulate critical processes of cell growth, death, EMT, metastasis, chemoresistance, radio-resistance, and immune-resistance in cancers. However, the role of moonlighting proteins (MLPs) in these actions of curcumin and resveratrol are unclear. We explain how curcumin and resveratrol control a complex self-regulated network of human MLPs in cancers. Both phytochemicals have similar modes of regulating oncogenic, regulatory, and tumour suppressive MLPs, and also have coordinated regulation of important MLPs (CTNNB1, MMP2, p53) and their activators and inhibitors. Notably, curcumin and resveratrol tightly control wild type p53 expression and activity by regulating activators, inhibitors, and gene targets of p53. Accordingly, curcumin and resveratrol can have anti-cancer effects in cancer cells with wild type or mutant p53. Overall, these mechanisms add insights into synergistic anti-cancer mechanisms of curcumin and resveratrol. We also identify novel mechanisms underlying pleiotropic / ‘off-target’ effects of curcumin and resveratrol. Such pleiotropic effects can arise when both phytochemicals regulate MLPs with opposing main and moonlighting functions, and MLPs with main and moonlighting functions in different cellular locations.

## Acknowledgements

Authors thank Dr. Rama Vaidyanathan and Dr. Reena Das of Dr. A.P.J. Abdul Kalam Centre (Dr. M.G.R. Educational and Research Institute) for their support.

## REFERENCES

1. Patel SS, Acharya A, Ray RS, Agrawal R, Ramsaneh R, Jain P. Cellular and molecular mechanisms of curcumin in prevention and treatment of disease. Crit Rev Food Sci Nutr. 60(6):887–939. 2020. DOI: 10.1080/10408398.2018.1552244.

2. Giordano A, Tommonaro G. Curcumin and Cancer. Review Nutrients 11(10):2376. 2019. DOI: 10.3390/nu11102376

3. Sferrazza G, Corti M, Brusotti G, Pierimarchi P, Temporini C, Serafino A, Calleri E. Nature-derived compounds modulating Wnt/ β -catenin pathway: a preventive and therapeutic opportunity in neoplastic diseases. Acta Pharm Sin B. 10 (10):1814–1834. 2020. DOI: 10.1016/j.apsb.2019.12.019

4. Ghoran SH, Calcaterra A, Abbasi M, Taktaz F, Nieselt K, Esmaeil Babaei E. Curcumin-Based Nanoformulations: A Promising Adjuvant towards Cancer Treatment. Molecules. 2022 Aug 16;27(16):5236. DOI: 10.3390/molecules27165236.

5. Franco-Serrano L, Hernández S, Calvo A, Severi MA, Ferragut G, Pérez-PonsJA 1, Jaume Piñol J, Pich O, Mozo-Villarias A, Amela I, et al. Querol E, Cedano J MultitaskProtDB-II: an update of a database of multitasking/moonlighting proteins. Nucleic Acids Res 4;46(D1):D645–D648. 2018. DOI:10.1093/nar/gkx1066

6. http://wallace.uab.es/multitaskII/proteins_list.php?a=integrated&ctlSearchFor=Homo+sapiens&simpleSrchFieldsComboOpt=&simpleSrchTypeComboNot=&simpleSrchTypeComboOpt=&criteria=and

7. Lo-Surdo P, Iannuccelli M, Contino S, Castagnoli L, Licata L, Cesareni G, Perfetto L. SIGNOR 3.0, the SIGnaling network open resource 3.0: 2022 update. Nucleic Acids Res. 2022. DOI: 10.1093/nar/gkac883.

8. Lorena A-C, Majano P, Sánchez-Toméro JA, Selgas R, López-Cabrera, Aguilera A, Mateo GG. Natural Plants Compounds as Modulators of Epithelial-to-Mesenchymal Transition. Front Pharmacol. 10:715. 2019. DOI: 10.3389/fphar.2019.00715.

9. Singh A, Settleman J. EMT, cancer stem cells and drug resistance: an emerging axis of evil in the war on cancer. Oncogene. 29(34):4741–51. 2010. DOI: 10.1038/onc.2010.215

10. https://www.genecards.org/cgibin/carddisp.pl?gene=CTNNB1&keywords=ctnnb1

11. Wong SHW, Fang CM 2, Chuah L-H 3, Leong CO, and Ngai SC: E-cadherin: Its dysregulation in carcinogenesis and clinical implications. Crit Rev Oncol Hematol. 121:11–22. 2018. DOI.org/10.1016/j.critrevonc.2017.11.010

12. Ferber EC, Kajita M, Wadlow A, Tobiansky L, Niessen C, Ariga H, Daniel J, and Fujita Y: A role for the cleaved cytoplasmic domain of E-cadherin in the nucleus. J Biol Chem. 283(19):12691–700.2008. DOI: 10.1074/jbc.M708887200

13. https://www.genecards.org/cgibin/carddisp.pl?gene=GSK3B&keywords=gsk3b

14. Ventura R, Kasia Mordec K, Waszczuk J, Wang Z, Lai J, Fridlib M, Buckley D, Kemble G, and Heuer TS: Inhibition of de novo Palmitate Synthesis by Fatty Acid Synthase Induces Apoptosis in Tumor Cells by Remodeling Cell Membranes, Inhibiting Signaling Pathways, and Reprogramming Gene Expression. EBioMedicine. 2(8):808–24. 2015. DOI: 10.1016/j.ebiom.2015.06.020

15. https://www.genecards.org/Search/Keyword?queryString=hmgb1

16. Adamo A, Frusteri C, Pallotta MT, Pirali T, Sartoris S, Ugel S. Moonlighting Proteins Are Important Players in Cancer Immunology. Review Front Immunol . 11:613069. 2021. DOI: 10.3389/fimmu.2020.613069

17. Grumolato L, Liu G, Haremaki T, Mungamuri SK, Phyllus Mong P, Akiri G, Lopez-Bergami P, Arita A, Anouar Y, Mlodzik M et al, β-Catenin-independent activation of TCF1/LEF1 in human hematopoietic tumor cells through interaction with ATF2 transcription factors. PLoS Genet. 2013;9(8):e1003603. DOI: 10.1371/journal.pgen.1003603.

18. https://www.genecards.org/cgi-bin/carddisp.pl?gene=ATF2&keywords=atf2

19. Ten Dijke P, Goumans M-J, Itoh F, and Itoh S: Regulation of cell proliferation by Smad proteins. J Cell Physiol. 191(1):1–16. 2002. DOI: 10.1002/jcp.10066

20. Mu Y, Sundar R, Thakur N, Ekman M, Gudey SK, Yakymovych M, Hermansson A, Dimitriou H, Bengoechea-Alonso MT, Ericsson J, Heldin CH, Landstrom M. TRAF6 ubiquitinates TGF [beta] type I receptor to promote its cleavage and nuclear translocation in cancer. Nat Commun. 2011; 2:330. DOI: 10.1038/ncomms1332

21. Seong H-A, Jung H, Kim K-T, Ha H. 3-Phosphoinositide-dependent PDK1 negatively regulates transforming growth factor-beta-induced signaling in a kinase-dependent manner through physical interaction with Smad proteins. J Biol Chem. 282(16):12272–89. 2007. DOI: 10.1074/jbc.M609279200.

22. https://www.genecards.org/cgibin/carddisp.pl?gene=SMAD4&keywords=smad4

23. Fenderet AW, Nutter JM, Fitzgerald TL, Bertrand FE, Sigounas G. NOTCH-1 promotes stemness and epithelial to mesenchymal transition in colorectal cancer. J Cell Biochem. 116(11):2517–27, 2015. DOI: 10.1002/jcb.25196

24. Schnell SA, Ambesi-Impiombato A, Sanchez-Martin M, Belver L, Xu L, Qin Y, Ryoichiro Kageyama R, Adolfo A Ferrando AA: Therapeutic targeting of HES1 transcriptional programs in T-ALL. Blood. 30;125(18):2806–14. 2015. DOI: 10.1182/blood-2014-10-608448

25. Tian C, Yu Y, Jia Y, Zhu L, Zhang Y: HES1 activation suppresses proliferation of leukemia cells in acute myeloid leukemia. Ann Hematol 94(9):1477–83. 2015. DOI: 10.1007/s00277-015-2413-0

26. Wang H, Li J, He J, Liu Y, Feng W, Zhou H, Wei H, Lu Y, Wanxin Peng P, Du F et al, Methyl-CpG-binding protein 2 drives the Furin/TGF-β1/Smad axis to promote epithelial-mesenchymal transition in pancreatic cancer cells. Oncogenesis. 9(8):76. 2020. DOI: 10.1038/s41389-020-00258-y

27. Zhu F, Wu Q, Ni Z, Lei C, Li T, Shi Y: miR-19a/b and MeCP2 repress reciprocally to regulate multidrug resistance in gastric cancer cells. Int J Mol Med. 42(1):228–236. 2018. DOI: 10.3892/ijmm.2018.3581

28. https://www.genecards.org/cgibin/carddisp.pl?gene=MMP2&keywords=mmp2

29. Horak CE, Lee JH, Elkahloun AG, Boissan M, Dumont S, Maga TK, Arnaud-Dabernat S, Palmieri D, Stetler-Stevenson WG, Lacombe ML, et al Paul S Meltzer, Patricia S Steeg. Nm23-H1 suppresses tumor cell motility by down-regulating the lysophosphatidic acid receptor EDG2. Cancer Res.67(15):7238–46. 2007. DOI: 10.1158/0008-5472.CAN-07-0962.

30. https://www.genecards.org/cgi-bin/carddisp.pl?gene=NME1&keywords=nme1

31. https://www.genecards.org/cgi-bin/carddisp.pl?gene=TIMP1&keywords=timp1

32. Cooper LC, Prinsloo E, Edkins AL, Blatch GL: Hsp90α/β associates with the GSK3β/axin1/phospho-β-catenin complex in the human MCF-7 epithelial breast cancer model. Biochem Biophys Res Commun. 413(4):550–4. 2011. DOI: 10.1016/j.bbrc.2011.08.136

33. Burrows F: Zhang H, Burrows F: Targeting multiple signal transduction pathways through inhibition of Hsp90. Review J Mol Med (Berl).82(8):488–99. 2004. DOI: 10.1007/s00109-004-0549-9

34. Nagaraju GP, Zhu S, Ko JE, Ashritha N, Kandimalla R, Snyder JP, Shoji M, Bassel F, El-Rayes. Antiangiogenic effects of a novel synthetic curcumin analogue in pancreatic cancer. Cancer Lett. 357(2):557–65. 2015. DOI: 10.1016/j.canlet.2014.12.007

35. Huang W-C, Huang W-C, Chen Y-J, Li L-Y, Wei Y-L, Hsu S-C, Tsai S-T, Chiu P-C, Huang W-P, Wang Y-N, Chen C-H, et al. Nuclear translocation of epidermal growth factor receptor by Akt-dependent phosphorylation enhances breast cancer-resistant protein expression in gefitinib-resistant cells. J Biol Chem. 286(23):20558–68. 2011. DOI: 10.1074/jbc.M111.240796

36. Li S, Payne S, Wang F 3, Claus P 4, Su Z 5, Groth J, Geradts J, de Ridder G, Alvarez R, Marcom PK, et al: Nuclear basic fibroblast growth factor regulates triple-negative breast cancer chemo-resistance. Breast Cancer Res. 17(1):91. 2015. DOI: 10.1186/s13058-015-0590-3

37. van Helvoort A, Smith AJ, Sprong H, Fritzsche I, Schinkel AH, Borst P, van Meer G. MDR1 P-glycoprotein is a lipid translocase of broad specificity, while MDR3 P-glycoprotein specifically translocates phosphatidylcholine. Cell. 87(3):507–17. 1996. DOI: 10.1016/S0092-8674(00)81370-7

38. https://www.genecards.org/cgi-bin/carddisp.pl?gene=CYCS&keywords=cycs

39. Chipuk JE, Kuwana T, Bouchier-Hayes L, Droin NM, Newmeyer DD, Schuler M, Green DR: Direct activation of Bax by p53 mediates mitochondrial membrane permeabilization and apoptosis. Science 303(5660): 1010–1014.2004. DOI: 10.1126/science.1092734

40. https://www.genecards.org/cgi-bin/carddisp.pl?gene=MDM2&keywords=mdm2

41. Rizki A, Mott JD, Bissell MJ. Polo-like kinase 1 is involved in invasion through extracellular matrix. Cancer Res.67(23):11106–10, 2007. DOI: 10.1158/0008-5472.CAN-07-2348.

42. Kobayashi EH, Kobayashi EH, Suzuki T, Funayama R, Nagashima T, Hayashi M, Sekine H, Tanaka N, Moriguchi T, Motohashi H, Nakayama K, et al: Nrf2 suppresses macrophage inflammatory response by blocking proinflammatory cytokine transcription. Nat Commun 23(7): 11624. 2016. DOI: 10.1038/ncomms11624

43. https://www.genecards.org/cgi-bin/carddisp.pl?gene=NQO1&keywords=nqo1

44. Narayanan 1, Nargi D, Randolph C, Narayanan BA. Liposome encapsulation of curcumin and resveratrol in combination reduces prostate cancer incidence in PTEN knockout mice. Int J Cancer. 125(1):1–8. 2009. DOI: 10.1002/ijc.24336

45. González-Sarrías A, Iglesias-Aguirre CE, Cortés-Martín A, Vallejo F, Cattivelli A, Pozo-Acebo LD, Del Saz A, de Las Hazas MCL, Dávalos A, Espín JC. Milk-Derived Exosomes as Nanocarriers to Deliver Curcumin and Resveratrol in Breast Tissue and Enhance Their Anticancer Activity. Int J Mol Sci. 23(5):2860. 2022. DOI: 10.3390/ijms23052860

46. Min K-W, Lee S-H, Baek SJ, Moonlighting proteins in cancer. Cancer Lett. 370(1):108–16. 2016. DOI: 10.1016/j.canlet.2015.09.022.

