## Supplementary Table 2 for "Analysis of Moonlighting Proteins Regulated by Curcumin and Resveratrol in Cancer"

| **Interacting**  **MLPs** | **Score** | **Outcome of MLP Interaction** | **PubMed**  **Number**  **(PMID)** |
| --- | --- | --- | --- |
|  |  | **GROUP 1 MLPs** |  |
| **CDH1 and CTNNB1 *** | SIGNOR = 0.963  STRING = 0.999 | CDH1 and CTNNB1 Complexes maintain adhesion of epithelial cells | **2927906** |
| **GSK3β and CDH1** | SIGNOR = 0.555  STRING = 0.606 | GSK3β phosphorylates CDH1. This strengthens CDH1-CTNNB1 binding. | 10671552 |
| **GSK3β and CTNNB1** | SIGNOR = 0.840  STRING = 0.968 | GSK3β phosphorylates CTNNB1 and triggers its degradation | 22083140 |
| **P53- HMGB1 *** | SIGNOR = NA  STRING = 0.951 | TP53-HMGB1 Complexes regulate apoptosis and autophagy | 22647615 |
| **GSK3β and SMAD3** | SIGNOR = 0.492  STRING = 0.521 | GSK3β phosphorylates SMAD3 and triggers its degradation*.* | **18172167** |
| **SMADs 3,4 and ATF2** | SIGNOR = 0.593  STRING = 0.285 | *SMADs 3 and 4 bind and increase transcriptional activity of ATF2* | **29665790** |
| **NOTCH1 and HES1** | SIGNOR = 0.745  STRING = 0.387 | *NOTCH1 Upregulates HES1* | **29665790** |
| **NOTCH1 and SMAD3** | SIGNOR =0.60  STRING = 0.285 | *Increases TGFB1 dependent HES1 expression. Promotes EMT* | 1463885735457181 |
| **NOTCH1 and**  **CTNNB1** | SIGNOR = 0.749  STRING = 0.400 | *NOTCH1 Activates Canonical Wnt*  *NOTCH1 Inhibits Canonical Wnt*  *CTNNB1 Regulates NOTCH1 activity* | 28947978  33907274  19000719 |
| **NOTCH1 and** **MDM2** | SIGNOR = 0.470  STRING = 0.486 | MDM2 ubiquitinates and activates intracellular domain of NOTCH1 | 23252402 |
| **NME1 and MMP2** | SIGNOR =0.307  STRING = NA | *NME1 Downregulates MMP2 Gene* | 17671192 |
|  |  | **GROUP 2 AND 3 MLPs** |  |
| **Hsp90AA1 and**  **TGFβR1** | SIGNOR =0.438  STRING = 0.715 | HSP90 Stabilizes TGFβRI  HSP90 -TGFβRI Interact in Fibrosis | 18591668  **27418101** |
| **PLK1** **and MDM2** | SCORE =0.462  STRING = NA | PLK1 phosphorylates and stimulates MDM2-mediated turnover of P53 | 19833129 |
| **P53 and MMP2** | SIGNOR = 0.401  STRING = NA | *P53 Upregulates MMP2 Gene* | **26245666** 10029407 |
| **P53 and CRYAB** | SIGNOR = 0.466  STRING = 0.486 | *CRYAB binds P53. P53 Upregulates CRYAB Gene* | **19799611** |
| **P53 and CYCS** | SIGNOR = 0.411  STRING = NA | *P53 Upregulates CYCS Gene* | 19007744 |
| **P53 and EGFR** | SIGNOR =0.54  STRING = 0.279 | *P53 Activates EGFR promoter and Upregulates EGFR Gene* | **8887630**  10029407 |
| **P53 and FGF2** | SIGNOR =0.396  STRING = NA | *P53 Downregulates FGF2 Gene* | 11313915  **11498784** |
| **P53 and MDR1** | SIGNOR =0.565  STRING = NA | *P53 Downregulates MDR1 Gene* | **11920581 29416681** |
| **NRF2 and NQO1** | SIGNOR = 0.435  STRING = NA | *NRF2 Upregulates NQO1 Gene* | 24024136 |
| **P53 and NQO1** | SIGNOR = NA  STRING = 0.835 | NQO1 Protects P53 from Proteasomal Degradation | **15687255** |

**Supplementary Table 2: Interactions between MLPs Regulated by Curcumin and Resveratrol.** MLPs which form a Complex (*). Most Interactions controlling Stability and Activity have SIGNOR and STRING scores (Underlined). Most Interactions controlling Gene Transcription / Expression only have SIGNOR scores (*Italics*). Not available (NA).
