## Supplementary Table 3 for "Analysis of Moonlighting Proteins Regulated by Curcumin and Resveratrol in Cancer"

| **Drug And MLP** | **Cell /Animal Model** | **Mechanism Of Curcumin OR Resveratrol Action** | **Effect of Curcumin OR Resveratrol** | **PMID Number** |
| --- | --- | --- | --- | --- |
| **Curcumin**  **CTNNB1** | Breast Cancer Stem Cells  Colon Cancer Line  Lung Cancer Line, Breast Cancer Line. | Inhibits CTNNB1 Nuclear Translocation and Activity  Same As Above  Downregulates CTNNB1 | Induces Apoptosis  Induces Apoptosis  Inhibits Growth, Invasion | **25315241**  **31584928**  **3046849831005718** |
| **Resveratrol**  **CTNNB1** | Osteosarcoma line  Colon Cancer Line  And Xenografts.  Uterine Cancer Line | Downregulates CTNNB1  Same As Above  Same As Above | Reduces Cell Invasion  Inhibits Growth  Induces Apoptosis | 29340064  2293544724756222  30867708 |
| **Curcumin**  **GSK3β** | Breast Cancer Line, Human Embryonic Carcinoma  Mouse Colon Carcinogenesis  Human Breast Cancer Line. | Activates GSK3β  Activates GSK3β  Downregulates GSK3β | Induces Apoptosis  Chemoprevention  Inhibits Growth, Invasion. Induces Death | 2013882925755051  21887819  36003508 |
| **Resveratrol**  **GSK3β** | Human Ovarian And Pancreatic Cancer Line, Lesions  Acute Lymphoblastic Leukaemia | Inactivates Nuclear GSK3β  Activates GSK3β | Induces Growth Arrest Decreases Proliferation  Induces Apoptosis | 2223458326556864  17868649. |
| **Curcumin CHD1** | Cancer Stem Cells  Colorectal Tumors  Nasal Cancer Xenografts | Increases CDH1 and CDH1-CTNNB1 Complexes.  Upregulates CDH1  Upregulates CDH1 | Inhibits EMT  Inhibits EMT  Stops Cell Migration | 25315241  23970932  20683022 |
| **Resveratrol**  **CHD1** | Colon Cancer Line  Lung And Bladder Cancer Line | Upregulates CDH1.  Upregulates CDH1 | Inhibits Cell Migration  Inhibits Cell Migration | 25884904  2314676036441214 |
| **Curcumin**  **FASN** | Drug Resistant Mouse Liver Cancer  Primary, Metastatic Colon Cancer Lines | Downregulates  FASN  Same As Above | Decreases Lipid Synthesis  Decreases Cell Migration | 32613952  25451013 |
| **Resveratrol**  **FASN** | Her2 Positive Breast Cancer Line  Review on Natural Products. | Downregulates FASN, HER2  Inhibits FASN Activity | Induces Apoptosis  FASN Specific Inhibition | 25448084  34459114 |
| **Curcumin**  **HMGB1** | Human malignant mesothelioma cells  Gastric Cancer lines  Review | Increases HMGB1 Secretion  Downregulates HMGB1  Decreases HMGB1 Levels and Release | Induces Pyroptosis  Induces Apoptosis  Prevents Sepsis | 24431405  31266378  20969478 |
| **Resveratrol**  **HMGB1** | Human Ovarian  Cancer line  Lewis Lung  Carcinoma  Rat Liver | Increases HMGB1 Secretion  Downregulates HMGB1  Downregulates HMGB1 | Induces Pyroptosis  Decreases Immuno-resistance  Prevents Injury | 31636696  29959821  31041919 |
| **Curcumin**  **ATF2** | Not Available |  |  |  |
| **Resveratrol**  **ATF2** | Benzopyrene [BP] induces Cell Death  Kidney, Hepatoma  Cell Lines | Inhibits ATF2, and reverses BP induced damage  Stimulates ATF2 and its Target Genes | Cytoprotective in Rat  Tumor Suppressive: | 27162022  26446263 |
| **Curcumin**  **TGFβR1** | Rat Hepatic Cells  Renal Epithelium | Downregulates TGFβRI  Same As Above | Induces Apoptosis  Inhibits EMT, Fibrosis | 17531121  23544048 |
| **Resveratrol**  **TGFβR1** | Not Available |  |  |  |
| **Curcumin**  **SMAD2,3** | Rats  Human Renal Epithelium | Decreases Expression of Phospho-Smad2/3  Same As Above | Decreases Fibrosis  Decreases Fibrosis | 22823335  34751624 |
| **Resveratrol**  **SMAD2,3** | Human Breast Cancer Xenografts  Rat Fibrosis | Downregulates SMADs 2,3  Inhibits SMADs 2,3 Phosphorylation | Inhibits Cell Migration  Prevents Fibrosis | 30901941  32952652 |
| **Curcumin**  **SMAD4** | Osteosarcoma Line  Human Fibroblasts | Inhibits SMAD4 Expression  Downregulates SMAD4 | Inhibits Proliferation  Inhibits Proliferation and Fibrosis | 26823826  33755170 |
| **Resveratrol**  **SMAD4** | Human Alveolar Rhabdomyosarcoma  Human Fibroblasts | Inhibits SMAD4 Expression  Downregulates SMADs 2,3,4 | Decreases Proliferation  Induces Apoptosis | 27008654  25721869 |
| **Curcumin NOTCH1**  **HES1** | Oesophageal and  Cholangiocarcinoma Cell Lines  Melanoma Cell Lines | Downregulates NOTCH1, HES-1  Downregulates NOTCH1, HES-1 | Induces Apoptosis,  Inhibits proliferation and invasion of cells. | 22363450  25890434 |
| **Resveratrol NOTCH1** | Ovarian, Breast Cancer Lines  Glioblastoma lines  Thyroid Cancer lines | Inhibits NOTCH1 signalling  Induces NOTCH1 signalling  Induces NOTCH1 signalling | Induces Apoptosis,  Induces Cytotoxicity  Inhibits Proliferation | 3081651332745729  21743969  23594881 |
| **Curcumin**  **MECP2** | Rat Hepatic Cells  Breast Cancer Tissue | Downregulates MECP2  Same As Above | Reduces Loss of Cells  Chemoprevention | 30367912  23593078 |
| **Resveratrol**  **MECP2** | Breast Cancer Tissue  Human Breast  Cancer lines | Downregulates MECP2  Same As Above | Chemoprevention  Induces Apoptosis | 23593078  31173924 |
| **Curcumin**  **MMP2** | Lung Cancer cells and Xenografts.  Prostate Cancer Cells and Tumors in Mice.  Human Leukaemia Cells in Mouse | Downregulates MMP2  Decreases MMP2 expression  Decreases MMP2 expression | Inhibits Invasion, Metastasis  Inhibits Growth, Invasion  Inhibits Growth, Invasion | 24445042  16389264  **31854220** |
| **Resveratrol**  **MMP2** | Hepatocarcinoma Lines  Lung Cancer Line | Inhibits MMP2  Downregulates MMP2. | Decreases Cell Invasion  Sensitizes to Taxol Induced Apoptosis | 20131808  28883751 |
| **Curcumin**  **TIMP1** | Pancreatic Cancer Lines  Nasopharyngeal Cancer Xenografts | Upregulates TIMP1,  Upregulates TIMP1 | Suppresses Metastasis  Inhibits Tumor Formation | 32945513  24482695 |
| **Resveratrol**  **TIMP1** | Hepatocarcinoma Lines  Lung Cancer Line | Increases TIMP1  Upregulates TIMP1 | Decreases Invasion  Sensitizes to Taxol Induced Apoptosis | 20131808  28883751 |
| **Curcumin**  **NME1** | B16F10 Mouse Adhesion Assay and Lung Metastases  Mouse Tumors | Increases Expression of NME1 and CDH1  Same As Above | Decreases Metastasis  Inhibits Metastasis. Increases life span | **12678405**  **17569210** |
| **Resveratrol**  **NME1** | Not Available |  |  |  |

**Supplementary Table 3: Group 1 MLPs Regulated by Curcumin and Resveratrol.** Model System, Regulatory Mechanisms, and Effects of Curcumin and Resveratrol are shown.
