## Supplementary Table 4 for "Analysis of Moonlighting Proteins Regulated by Curcumin and Resveratrol in Cancer"

| **Drug And MLP** | **Cell /Animal Model** | **Mechanism Of Curcumin OR Resveratrol Action** | **Effect of Curcumin OR Resveratrol** | **PMID Number** |
| --- | --- | --- | --- | --- |
| **Curcumin**  **Hsp90 AA1** | Epidermoid Cancer  Pancreatic Cancer | Downregulates Hsp90  Same As Above | Induces Apoptosis  Anti-Angiogenic | 20039095  25497868 |
| **Curcumin CRYAB** | Astrocytoma cells.  Rats, Rat Eye | Induces CRYAB  Prevents CRYAB Downregulation | Neuroprotective  Prevents Toxicity, Cataract Formation | 22705585  **3310701821311744** |
| **Resveratrol**  **NOS3** | Endothelial  Progenitor Cell  Rats | Upregulates and Activates NOS3  Prevents NOS3 Downregulation | Produces Nitric Oxide  Prevents Vascular Damage | 25302702  29973698 |
| **Curcumin COX-1 / PTGS1** | Curcumin And Analogues  Rat Colon Carcinogenesis | Selective COX-1 Inhibitor  Same As Above | Anti-Inflammatory  Inhibits Tumor Intitation, Progression | 17201156  11748382 |
| **Resveratrol COX-1 / PTGS1** | COX-1 and 2 Inhibition Assays  Same As Above | Selective COX-1 Inhibitor  Inhibition of COX-1,2 | Anti-Inflammatory  Anti-Inflammatory | 15020596  15364641 |
| **Curcumin**  **EGFR** | Breast Cancers  Lung, Other Cancers | Downregulates EGFR  EGFR Inhibitor | Inhibits Tumor Growth  Same As Above | 18384098  2248919217569214. |
| **Resveratrol**  **EGFR** | Cervical cancer Line  Many Cancer Types | Downregulates EGFR  Inactivates EGFR | Inhibits Tumor Growth  Same As Above | 36558430  12840221 |
| **Curcumin**  **FGF2**    **Resveratrol**  **FGF2** | Not Available  Bladder Cancer Cells, Xenografts.  Fibrosarcomas In Chick Embryo  Endothelium | Downregulates FGF2  Inhibits FGF2-induced Angiogenesis, Growth  Same As Above | Induces Apoptosis and Inhibits Growth  Inhibits Growth  Suppresses Growth, Vascularization | 20028382  16091005  16675471 |
| **Curcumin**  **MDR1 / ABCB1** | Multidrug Resistant Osteosarcoma cells, Xenografts And Human Cervical cancer Line | Inhibits MDR1 Activity | Increases Drug Sensitivity  Inhibits Metastasis | 24229684  15476675 |
| **Resveratrol**  **MDR1 / ABCB1** | Drug Resistant Colorectal Cancer  Human Breast Cancer Xenografts | Downregulates MDR1 And / OR Inhibits MDR1 Activity  Same As Above | Reverses Multidrug Resistance  Increased Sensitivity to Doxorubicin | 26124005  24161697 |
| **Curcumin**  **CYCS** | Burkitt's Lymphoma  Human Breast and Prostate Cancer lines | Upregulates CYCS  Triggers CYCS Release | Induces Apoptosis  Induces Apoptosis | 19326074  15738001 |
| **Resveratrol**  **CYCS** | Ovarian Cancer Line  Human Transformed Breast Epithelium | Triggers CYCS Release  Same As Above | Induces Apoptosis  Induces Apoptosis | 1474478  26212257 |
| **Curcumin p53** | Liver Damage  Mouse Bladder Cancer.  Multiple Myeloma | Prevents p53 Decrease  Upregulates p53, p21  Upregulates p53 | Protects Liver  Decreases Growth, Increases Death  Inhibits Proliferation | **34231234.**  **22189739**  25789029 |
| **Resveratrol p53** | Lung Cancer Line  Mouse Skin Cancer Lines and Model  Mouse Epithelium | Upregulates p53  Upregulates p53  Destabilises P53 | Induces Apoptosis  Chemosensitizes Tumors to 5-FU  Increases Cell Survival | **32945383**  **25736303**  **28070015** |
| **Curcumin**  **MDM2** | Many Cancer Lines | Downregulates MDM2 | Decreases Growth OR Induces Apoptosis | 17332326**17900536**25789029 |
| **Resveratrol**  **MDM2** | Salivary Tumour  Lung Cancer Line  Squamous Carcinoma | Upregulates MDM2  Upregulates MDM2  Upregulates MDM2 | Induces Apoptosis  Induces Apoptosis  Increases Sensitivity to Cisplatin | 16081268  [23666059](https://pubmed.ncbi.nlm.nih.gov/23666059)  [34204834](https://pubmed.ncbi.nlm.nih.gov/34204834) |
| **Curcumin PLK1** | Human Cancer Lines | Downregulates PLK1 | Induces Growth Arrest and Apoptosis | 29501629 28239299 |
| **Resveratrol PLK1** | Human Breast Cancer Lines  Natural killer Cells,  T-Cell Lymphoma | Downregulates PLK1  Downregulates PLK1 | Inhibits Proliferation  Induces Apoptosis | 27109433  32617308 |
| **Curcumin NRF2** | 5-FU Resistant Colon Cancer line.  Human Oral Carcinoma  Rat Ovarian Cancer  Mouse Liver | Inhibits NRF2 Activity  Inhibits Translocation and Activity of NRF2  Induces NRF2.  Induces Translocation and Activity of NRF2 | Reverses Chemoresistance  Sensitizes to Cisplatin  Inhibits EMT, Fibrosis  Decreases Oxaliplatin Toxicity | 30365132  26469832  35517894  32021093 |
| **Resveratrol**  **NRF2** | Solid Tumors  Human Leukemia  Rat Mammary Gland  Mouse Colon Tumor | Inhibits or Down-regulates NRF2  Upregulates NRF2, NQO1  Activates NRF2 and its Gene Targets | Reverses Chemoresistance  Delays Estrogen Induced Tumors  Anti- Inflammatory Induces Apoptosis | 31802896  31032633  2489486  21355597 |
| **Curcumin NQO1** | Mouse Liver  Mouse Lymphoma | Induces Nuclear Translocation of NRF2  Activates NRF2, NQO1. Restores p53 | Decreases Oxaliplatin Toxicity  Anti-Cancer Effect | 32021093  25860911 |
| **Resveratrol**  **NQO1** | Rat Mammary gland.  Human Immortalized Breast Epithelial Line | Upregulates NRF2, NQO1  Upregulates NRF2, NQO1 and Reversed Estrogen Effects | Decreased Estrogen Induced Tumors  Estrogen decreased NRF2, NQO1 expression | 2489486  25130429 |

**Supplementary Table 4: Group 2 and 3 MLPs Regulated by Curcumin and Resveratrol.** Model System, Regulatory Mechanisms, and Effects of Curcumin and Resveratrol are shown. 5-Fluorouracil (5-FU)
